# Global conservation prioritization approach provides credible results at a regional scale

**DOI:** 10.1101/2023.08.31.555788

**Authors:** Michael Roswell, Anahí Espíndola

**Affiliations:** Department of Entomology, University of Maryland, College Park, MD, USA; Department of Biology, University of Maryland, College Park, MD, USA

**Keywords:** Predictive modelling, conservation status, random forest, lepidopterans, plants, downscaling, prioritization, multi-taxon

## Abstract

**Aim:** Conservationists and managers must direct resources and enact measures to protect species, despite uncertainty about their present status. One approach to covering the data gap is borrowing information from data-rich species or populations to guide decisions about data-poor ones, via machine learning. Recent efforts demonstrated proof-of-concept at the global scale, leaving unclear whether similar approaches are feasible at the local and regional scales at which conservation actions most typically occur. To address this gap, we tested a global-scale predictive approach at a regional scale, using two groups of taxa.

**Location:** State of Maryland, USA.

**Taxa:** Vascular land plants and lepidopterans.

**Methods:** Using publicly available occurrence and biogeographic data, we trained random forest classifiers to predict the State-level conservation status of species in each of the two focal taxa. We assessed model performance with cross-validation, and explored trends in the predictions.

**Results:** Our models had strong discriminatory ability, accurately predicting status for species with existing status assessments. They predict that the northwestern part of Maryland, USA, which overlaps the Appalachian Mountains, harbors a higher concentration of unassessed, but likely threatened plants and lepidopterans. Our predictions track known biogeographic patterns, and unassessed species predicted as most likely threatened in Maryland were often recognized as also needing conservation in nearby jurisdictions, providing external validation to our results.

**Main Conclusions:** We demonstrate that a modelling approach developed for global analysis can be downscaled and credible when applied at a regional scale that is smaller than typical species ranges. We identified select unassessed plant and lepidopteran species, and the western, montane region of Maryland as priority targets for additional monitoring, assessment, and conservation. By rapidly aggregating disparate data and integrating information across taxa, models like those we used can complement traditional assessment tools and assist in prioritization for formal assessments, as well as protection.

## Introduction

Conservationists and natural resource managers must direct resources and enact measures to assess – and ultimately protect – species and populations, despite uncertainty about their occurrence patterns, life history, demographic trends, or the specific threats they face. One stepping-stone approach to bridging the data gap is borrowing information from data-rich species or populations to guide decision making about data-poor ones using predictive modelling frameworks(Callaghan, Nakagawa, & Cornwell, 2021; Free et al., 2020; Pelletier, Carstens, Tank, Sullivan, & Espíndola, 2018; Zizka, Andermann, & Silvestro, 2022),. Increasingly, publicly available datasets that aggregate observations (Heberling, Miller, Noesgaard, Weingart, & Schigel, 2021) enable formulating and fitting predictive models to guide conservation and natural resource management (Bastin et al., 2019; Caetano et al., 2022; Callaghan et al., 2021; Panter et al., 2020; Pelletier et al., 2018; Zizka et al., 2022). The predictions from such models might be used to support conservation actions, and guide prioritization for resource-intensive status assessments (Pelletier et al., 2018). On the latter, national and subnational jurisdictions generate taxon-specific management plans triggered by formal status assessments (e.g., via National and State-level Endangered Species Acts in the United States (Department of the Interior, 1973; “Nongame and endangered species conservation act,” 2022)). In this context, status predictions from predictive models such as those mentioned above are not sufficient to trigger protections, but might undergird the listing process, by assisting with monitoring and assessment prioritization in a resource-limited world.

Several knowledge gaps or “shortfalls” prevent strategic species protection (Cardoso, Erwin, Borges, & New, 2011b; Mokany & Ferrier, 2011). Here, we focus on the “Rabinowitzian” shortfall, i.e., the gap between observing species’ occurrences, and using that information to evaluate their conservation risks (Cardoso, Erwin, Borges, & New, 2011a; Crisfield, Blanchet, & Gravel, 2022; Isbell et al., 2023; Rabinowitz, 1981). Models that use existing data can identify under-recognized patterns of threat and endangerment (Regan et al., 2008), and enable efficient allocation of monitoring, assessment, and protection resources (Hochkirch et al., 2021). For these goals, statistical learning tools that aggregate disparate data streams can recognize meaningful patterns that might escape notice in more granular analyses. Such tools were tested by Pelletier et al. (2018), who used only a handful of spatial and climatic covariates, and were able to predict the conservation status of unassessed land plants. They identified taxa likely to meet IUCN Red List criteria, as well as previously unrecognized hotspot regions that likely harbor high concentrations of threatened but unassessed taxa. The authors anticipated that their approach could serve conservation needs at smaller spatial scales, which would make their method directly applicable at the scales at which most conservation actions take place. The outputs from their models, which include predicted likelihood of threat for unassessed taxa, maps illustrating the occurrence patterns of species likely to be relatively threatened or secure, and a suite of covariates associated with the occurrence patterns of threatened species, provide decision makers with clues about the conservation needs of poorly known species. Because it is insufficient to focus conservation efforts solely on the needs of previously assessed species (Baker et al., 2019; Gallagher et al., 2023), these kinds of clues provide a starting place to expand the purview to unassessed taxa (Hochkirch et al., 2021). Furthermore, because Pelletier et al.’s (2018) predictive modeling approach uses existing data and can be iterated as additional information or priorities dictate, it is amenable to initial-path-setting decisions (e.g., whom to hire, where to monitor, which experts to engage) at local and regional scales (Pressey, Mills, Weeks, & Day, 2013).

Although the approach Pelletier et al. (2018) used can be technically implemented at nearly any scale, downscaling entails radical shifts in data availability, quality, and biogeographic representation that could improve or erode the utility of the approach. At smaller, regional scales, data quality may be more similar between assessed and unassessed taxa, which should improve predictions. Additionally, at smaller scales, complete datasets containing a wider variety of potentially appropriate covariates (e.g., land use and its history) may be available. Nevertheless, at smaller spatial scales, records of species’ occurrences are unlikely to cover species’ whole ranges, and the number of species in a smaller region is also likely to be smaller, potentially limiting statistical power. Additionally, it is unknown whether the approach is extensible to other taxa beyond plants without substantial modification. If the same suites of geographic covariates can predict conservation status of disparate taxa, this predictive approach has high promise to assist in a variety of conservation contexts.

Here, we downscale the approach Pelletier et al. (2018) took to predict conservation statuses, applying similar methods to vascular land plants (the focal higher taxon in Pelletier et al. (2018), hereafter “plants”) and lepidopterans (a new focal group) within a sub-national management region (the State of Maryland, USA). We selected these two taxa because the State of Maryland maintains a rare, threatened, and endangered list for each (Maryland Natural Heritage Program, 2021a, 2021b), and the ecology and conservation needs of these two taxa differ (Isbell et al., 2023; Wagner, Fox, Salcido, & Dyer, 2021). We asked whether, despite differences in the quality and quantity of data available, as well as the ecology of the taxa upon which predictions are made, a similar predictive modelling approach can help cover the Rabinowitzian shortfall, and guide conservation decisions at smaller, sub-national scales. We conducted this study as a test-of concept, relying on datasets that were publicly available and could be used with minimal processing. In discussing our results, we evaluate model performance and prediction credibility, and then explore our predictions. We identify species that are likely threatened in the State of Maryland, USA, and regions within the State that are likely to harbor a high density of threatened, unassessed species. Finally, we use our case studies to discuss the promises and potential pitfalls of using similar predictive modelling tools for conservation prioritization at regional and local scales.

## Materials and Methods

### Method Overview

To predict the conservation status of lepidopterans and plants in the state of Maryland, we obtained publicly available species occurrence and spatial data. We used data for all assessed species to build random forest classifiers, tuning hyperparameters and evaluating model discriminatory ability and expected generalization error through nested cross-validation. Then, for each taxon group (i.e., lepidopterans and plants), we fit a final model that was trained on all assessed species, and used these to predict conservation status for all unassessed species known to occur within the state. We mapped predictions to visually screen for spatial patterns, and then conducted a limited literature search to contextualize status predictions with independent lines of evidence. In the following sections, we provide details on the data we used, how we constructed and assessed models, the ways we generated and visualized predictions, and how we used independent lines of evidence to corroborate predictions.

### Data

#### Plant and Lepidopteran Records

We obtained observations of plants and lepidopterans from the State of Maryland from 2001-2023 from GBIF (Appendix S2), irrespective of life stage (GBIF.org, 2024). The timeframe matches that for the spatial covariates (see next section). We filtered the occurrence dataset to include only records associated with valid species observed in the wild, as discussed below. We assigned each species to a conservation status category, based on Maryland’s State-level rankings, as recorded in NatureServe (NatureServe, 2023). In order to do this, we first harmonized the taxonomies used by NatureServe and GBIF with programmatic tools from the R Packages *rgbif, natserv,* and *taxize* (Chamberlain, 2020; Chamberlain, Oldoni, et al., 2022; Chamberlain, Szoecs, et al., 2022), and some manual curation (Appendix S1), and then merged the NatureServe conservation status ranks with the occurrence data based on the consensus species identification. Because relatively few plants, butterflies, or moths had State conservation status in Maryland, we grouped their State conservation categories into three groups as follows: “secure” taxa were all those categorized as S4 or S5, “threatened” taxa were all those categorized as S1, S2, S3, SH, or SX, taxa categorized as SNA were removed, and all other taxa were considered unranked. To restrict our analysis to only species considered native to the State, we also filtered records based on the national and subnational exotic/native fields from NatureServe, and subsequently, for plants, also filtered them based on a recent Maryland flora (Knapp & Naczi, 2020). Three threatened, and 334 unranked lepidopteran species, and 91 threatened, 12 secure, and 59 unranked plant species were filtered out because they were represented by only a single occurrence location. Overall, our filtered dataset contained records for, respectively, 30, 90, and 1450 threatened, secure, and unranked lepidopterans, and 312, 134, and 952 threatened, secure, and unranked plant species. The 1570 lepidopteran species were represented by a total of 128,245 records; 1398 plant species were represented by 192,103 records. Additional species data, such as range characteristics or biological traits, are not available across all species in the study and therefore were not included as features in model constructions.

#### Environmental Variables

For each location record of a valid native species, we extracted latitude, longitude, and a suite of associated spatial variables. We considered spatial datasets that were complete across the study region, publicly available, and might plausibly be associated with species threat status. We tended to keep all variables from all datasets we could obtain since on the one hand, this was unlikely to worsen our predictions, as Random Forest models built on meaningful predictors tend to be robust to additional variables (Hastie, Tibshirani, & Friedman, 2017). On the other hand, doing so allowed us to evaluate the performance of this modeling approach in a generic context (i.e., predictions for a variety of taxa are all based on the same covariates). We extracted current and historic landcover, and indices of landcover change, from the National Land Cover Dataset (J. Wickham, Stehman, Sorenson, Gass, & Dewitz, 2021), which includes 30m x 30m landcover data at 5-year intervals from 2001 to 2016. We anticipated that these variables might be associated with threat *per se*, as well as different geomorphology. We also extracted interpolated climate data from the expanded CHELSA dataset, which includes 19 variables that summarize estimated daily, monthly, and annual temperature and precipitation (bioclim), and an additional 50 variables derived from physical data such irradiance, pressure, and humidity, as well as their relationships to biological tolerance ranges (e.g., growing season length estimates) at the resolution of 30 x 30 arc-seconds (Brun, Zimmermann, Hari, Pellissier, & Karger, 2022). Additionally, for each occurrence record, we extracted the estimated slope (at roughly 100m x 100m resolution) from the 1-m Statewide Slope layer (Eastern Shore Regional GIS Cooperative, 2021). We hypothesized that these variables might help distinguish geomorphology and microclimates within the study region, thereby enabling models to recognize unique niches and habitat types. Different spatial variables were available at different resolutions, and for each, we used the finest resolution available, extracting variable values at the coordinates of each occurrence record. In total, we included 69 spatial variables.

Although we gathered spatial data at the occurrence level, we fit models at the species level. To create the species-level datasets, we summarized each environmental variable across all occurrences of a species to a pair of summary statistics: for continuous variables we recorded the mean and standard deviation, and for categorical variables we recorded the modal value and the number of unique levels observed for the given species. In addition, for each species, we recorded the minimum and maximum latitude and longitude of their records. Variables with only a single value or missing values prevented model fitting and were thus dropped from the datasets for each group of organisms. To serve as a sort of null model for variable importance, we also included a random continuous variable as a predictor in each model. Models were built using these 153 summarized species-level predictors.

### Model Construction and Evaluation

We divided the dataset into a training dataset, comprised of all species within the categories “threatened” or “secure,” and a prediction dataset, comprised of all unranked species. In order to tune hyperparameters and evaluate model performance, while retaining as much training data as possible, we used nested repeated 10 x 10-fold cross-validation. To do this, we created 10 x 10 outer folds of the training dataset, and on each outer fold, trained a random forest classifier using 10 x 10-fold cross-validation, using the R packages *caret* and *randomForest* (Kuhn et al., 2022; Liaw, Wiener, Breiman, & Cutler, 2022). On the inner folds, we conducted a grid search over the hyperparameter “mtry,” which controls the number of features available to use for splitting the data at each node, and has been shown among the random forest hyperparameters to substantively affect model performance (Hastie et al., 2017; Probst, Wright, & Boulesteix, 2019). We left the other hyperparameters at their default values, as these are typically considered adequate for model construction (Probst et al., 2019). We used the area under the receiver-operating characteristic curve (AUROC) as our measure of performance in parameter tuning, as this measure emphasizes model discriminatory ability (Ling, Huang, & Zhang, 2003), rather than accuracy *per se*. This performance target should lead to models that can distinguish levels of threat (i.e., that rank species well), but does not hinge on a single choice about the costs of misclassifying threatened vs. relatively secure species. Although the Random Forest algorithm enables less computationally intensive model assessment using out-of-bag errors, here we used traditional cross-validation, which would be applicable to any model fitting algorithm (Kuhn et al., 2022).

For each of the 100 outer folds, we fit a “final model” with the entire dataset, save the held-out fold, based on the selected hyperparameters, and used this model to make predictions on the held-out fold. Using random forest votes as a measure of class probability, we computed AUROC, and to evaluate model performance on unseen data, we computed the mean and standard deviation of the 100 AUROC values computed on these 100 outer folds. In addition, we recorded the mean and standard deviation in AUROC from the inner, tuning folds, as large differences between AUROC from the inner and outer folds could indicate a lack of stability in model performance, or a tendency to overfit the training data.

After the extensive cross-validation described above, we re-trained each model (i.e., one for plants and one for lepidopterans) on the entire training dataset, again tuning the hyper-parameter “mtry” with 10x repeated 10-fold cross-validation. We used the “final model” for each taxon group from this step to make all predictions on unclassified species. The performance estimates we obtained from previous cross-validation provide the expected performance of these final models on unseen data, with two major caveats. First, additional data were used to train the final models, which are therefore not exactly equivalent to the models built on subsets of the same data. Second, we use these final models for predicting conservation status for unassessed, rather than assessed species. For unassessed species, we can only evaluate model performance indirectly, by comparing predictions with non-definitive external data (see below).

#### Optimal Threshold

Depending on the application, greater precaution may be warranted for misclassification errors for either threatened or relatively secure species, implying different ideal thresholds for the share of random forest “votes” required to classify a species into a certain category. As the threshold for classifying a species as threatened increases, fewer species will be classified as threatened, and the chance of mis-classifying a species that is, in fact, relatively secure as “threatened” falls, while the chance of mis-classifying a threatened species as “secure” rises. In evaluating model performance, we focused on discriminatory ability (using AUROC, which effectively considers all possible dichotomous thresholds (Hand, 2009)), and did not assess accuracy (i.e., the average rate of prediction errors) (Ling et al., 2003). However, in order to visualize predictions and more clearly discuss our results, we selected a dichotomous classification threshold, acknowledging that across different applications, other thresholds might be preferable (Walker, Leão, Bachman, Bolam, & Nic Lughadha, 2020). For each taxon group, we selected the threshold that maximized the geometric mean of sensitivity and specificity on out-of-fold training data using the function *performance* from the R package *ROCR* (Tobias, Sander, & Beerenwinkel, 2022).

#### Visualizing Predictions

We created state maps using the R packages *raster, tigris, sf, rnaturalearth,* and *ggplot2* (Etten et al., 2023; Massicotte, South, Hufkens, & Philippe, 2023; Pebesma et al., 2023; H. Wickham et al., 2022) to visualize model predictions and highlight areas in which many records of putatively threatened species were found. To visualize model predictions, we plotted each occurrence record using a color palette representing the random forest votes for “threatened” given to the taxon. To better visualize concentrations of threatened species, we divided the study region into 8km x 8km grid cells, and each cell value represented the number or proportion of species classified as threatened according to a threshold specific to each higher taxon (see “Optimal Threshold”, above). Such maps can be used to prioritize locations for additional survey and assessment resources, or to prioritize locations for conservation funding, as locations with a high concentration of putatively threatened species may both yield valuable data for evaluating species’ conservation statuses, and warrant precautionary protection.

#### Assessing Predictors

Random forest models are poorly suited to inference on causal relationships, but it is possible to diagnose which predictors most strongly drive model predictions. Towards this end, we measured “MDA” using the R package r*andomForest* (Bénard, Da Veiga, & Scornet, 2022; Breiman, 2001; Ishwaran, 2007; Liaw et al., 2022). MDA approximates the degree to which each variable improves the predictive accuracy of trees within the random forest. We ranked all variables based on their MDA scores within each cross-validation run such that the variable with the greatest MDA score in a particular run would be ranked first. We then plotted the distribution of MDA ranks across cross-validation runs for the top five predictors for each taxon group. Variables exhibiting consistent, low MDA ranks had relatively strong and/or clear associations with conservation status.

#### Literature Search

Although we cannot directly test whether predictions on unassessed species are correct without formal status assessments, we can use external data sources as additional lines of evidence to corroborate or challenge our predictions. To do this, we gathered external data for the 15 (or more, in case of ties) species for each taxon group that received the most and fewest votes for the category “threatened.” These species, for which the models had the strongest predictions, provide an opportunity to screen for inconsistencies that might suggest incorrect predictions, despite the fact that information on the conservation statuses of unassessed species is, by definition, lacking. For each of these species, we first asked if the species had been recognized as threatened in nearby jurisdictions by querying the species name on NatureServe (NatureServe, 2023)(Appendix S2) for subnational status assignments in Eastern North America (i.e., the continental USA east of the Mississippi, and the Canadian provinces of Ontario, Québec, New Brunswick, Nova Scotia, Prince Edward’s Island, and tribal/first nations lands within this region). A match between our status prediction and the recorded status in nearby locations supports our prediction. Second, we also examined contemporary geographic range data for the same top species. To do this, we visualized recent occurrence records in North America for each species using iNaturalist (iNaturalist Network, 2022). For species predicted as “threatened” (and the opposite for species predicted as “secure”), if it appeared that species current range limits may be in or near Maryland, we considered our predictions further supported.

## Results

### Model Construction and Performance

On average, the discriminatory abilities were high for models for both taxon groups (mean AUROC plants = 0.933, sd = 0.039; lepidopterans = 0.902, sd = 0.084; Fig. 1), with both models generally outputting higher “threatened” probabilities to threatened species than to secure ones (Fig. 2). We did not detect large differences in AUROC between the inner and outer folds for lepidoptera, but did for plants. This difference indicates less stable models for plants (i.e., model performance may be very sensitive to the inclusion of particular training data). On the other hand, hyperparameter selection was highly variable for lepidopteran and somewhat more consistent for plant models (mean mtry for plant models = 6.9, sd = 14.0, final model mtry = 2; lepidopterans = 61.0, sd = 60.0, final = 140). When we set a vote threshold for classification that maximized model sensitivity and specificity, the optimal vote threshold differed between the two taxon groups: 0.66 for plants and 0.36 for lepidopterans. At these thresholds, both models accurately classify previously assessed species.

**Figure 1.**
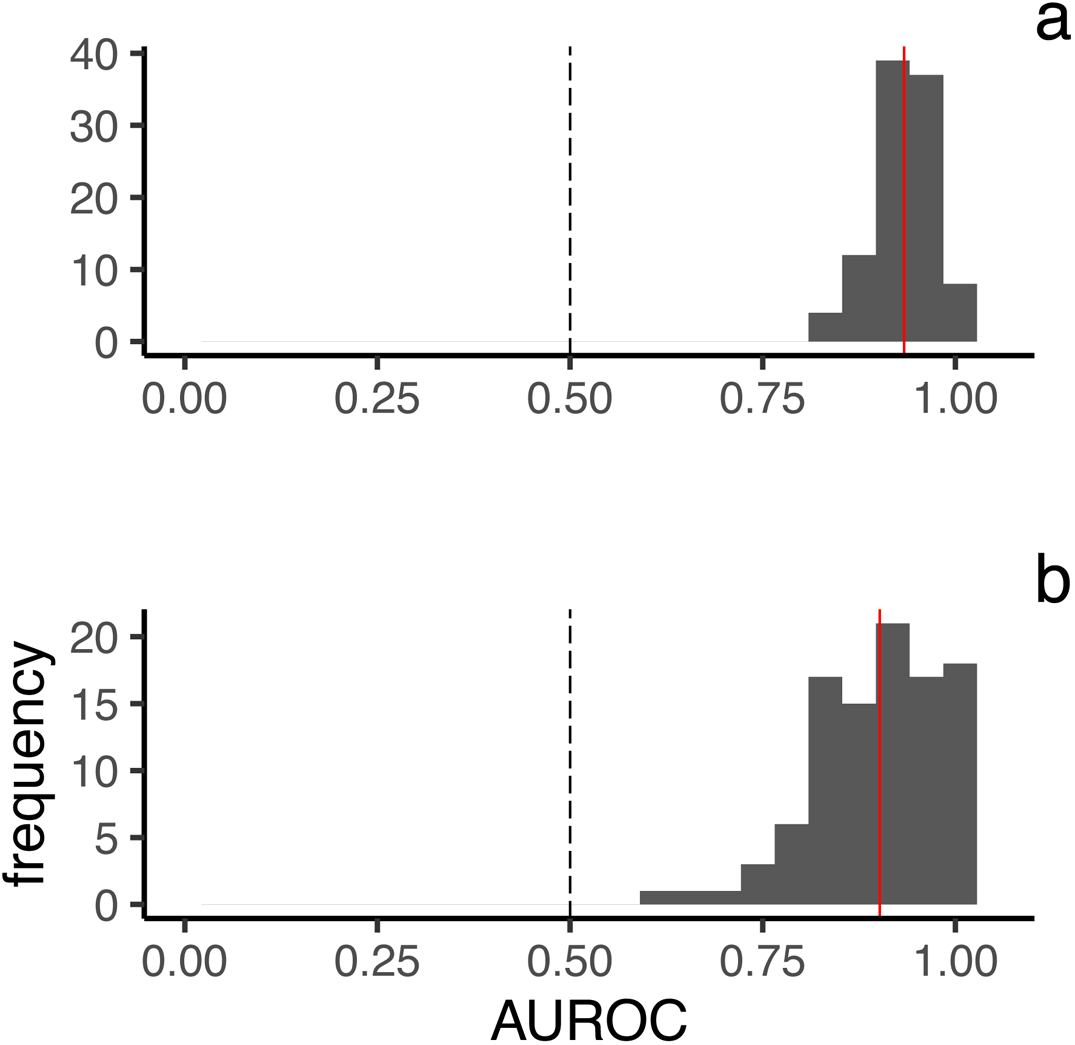
Discriminatory ability of random forest models was high for both plants (a) and lepidopterans (b). Histograms show distribution of area under the receiver operating curve (AUROC) values on unseen data in 10x repeated 10-fold cross-validation. In each panel, mean AUROC is shown with a red vertical line; dashed vertical line indicates the expected performance of a model making random guesses (AUROC of 0.50).

**Figure 2.**
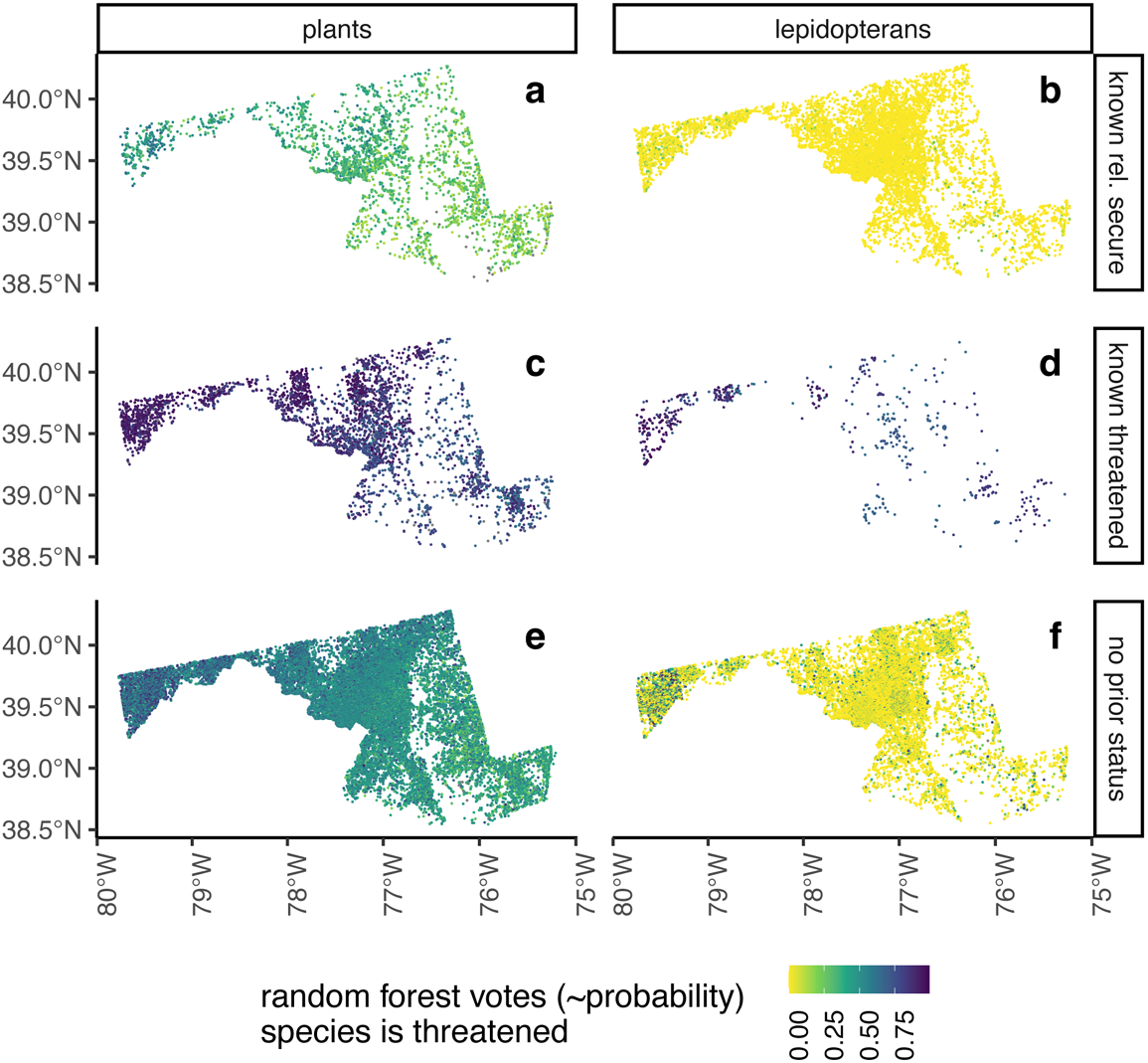
Random forest predictions (votes) from final models on occurrence (point) records of assessed (a-d) and unassessed (e, f) species. Models distinguished threatened and relatively secure species in the training dataset (lighter points in panels a and b, darker in c and d), and identify a concentration of putatively threatened, unassessed species in Western Maryland (trend towards darker points in western portion of maps in e, f). Color gradient indicates predicted values.

### Predictions for unassessed species

At the selected vote thresholds, our classifiers predicted 52% (493/952) of unassessed plant species and 44% (639/1450) of unassessed lepidopteran species as threatened in Maryland (Table S1). The most predictive variables differed between plants and lepidopterans (Fig. 3). For lepidopterans, variables related to coldest temperatures tended to place among the top variables. These included the mean first day of year above 5° C, and the latitudes and longitudes of Maryland records. For plants, the top variables were often associated with variability, such as that in land use change index, and in daily mean air temperature both during the coldest month and during the growing season, and also in daily temperature ranges. The mean first day over 10°C was also an important predictor for plant conservation status.

**Figure 3.**
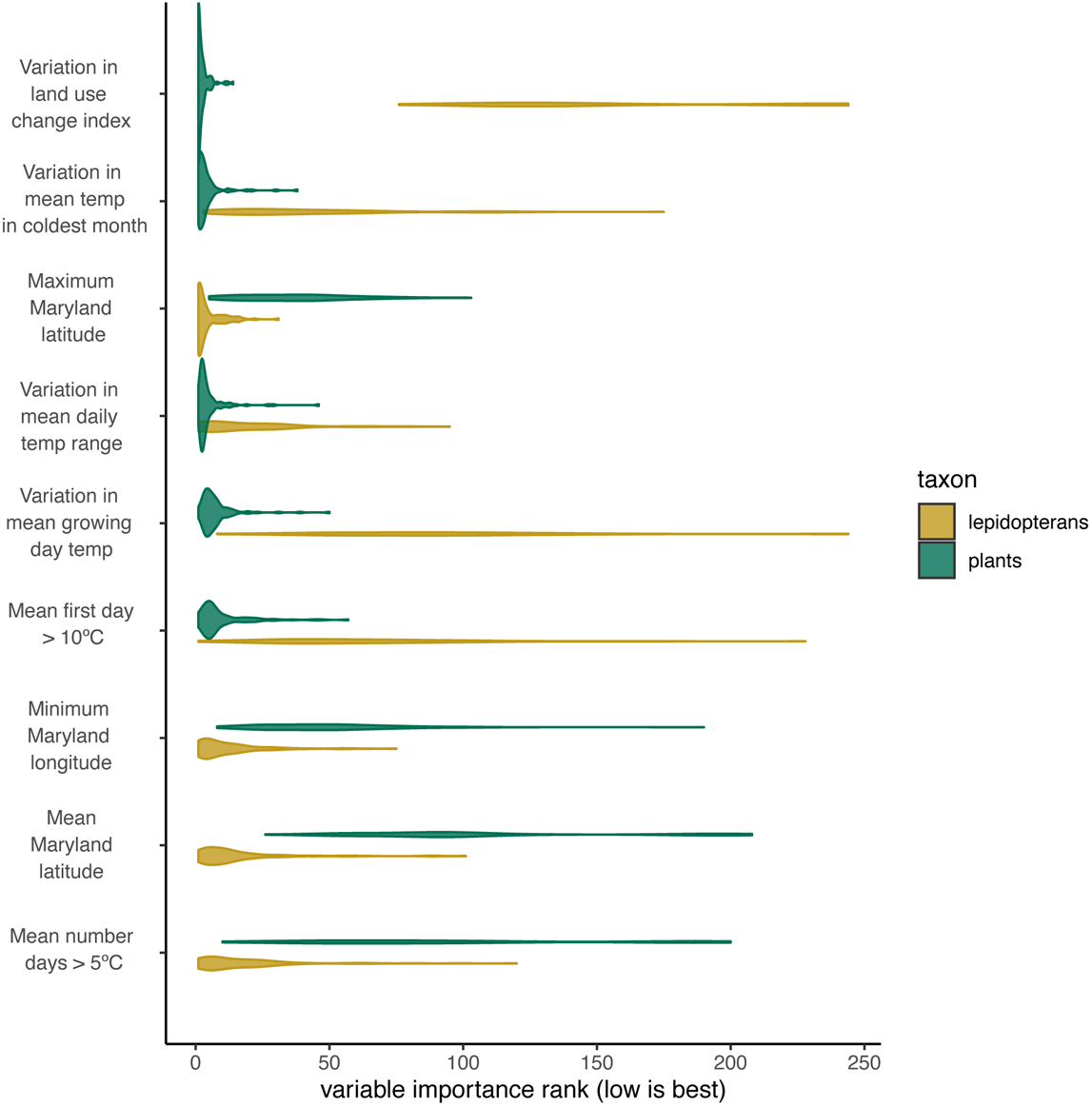
Distribution of variable importance ranks for the five predictors with the lowest mean MDA ranks (i.e., those with higher importance across cross-validation runs) for plants (green) and lepidopterans (yellow). Maximum ranks exceed the total number of variables because each level of categorical variables receives its own MDA measure.

Model predictions highlighted the northwesternmost part of the state as a possible conservation hotspot for both plants and lepidopterans (Fig. 4). Although unassessed species predicted as “threatened” were found throughout the state, their occurrence records appear concentrated in the northwestern half of Maryland (Fig. 4). Furthermore, the mean per-cell predicted threat probabilities showed a strong east-west gradient, with the highest mean threat probabilities in the westernmost parts of Maryland (Fig. S1). For both taxa, a higher number, but not high proportion, of species were predicted as “threatened” in the Piedmont region in the center of the state (Figs. 4, S1).

**Figure 4.**
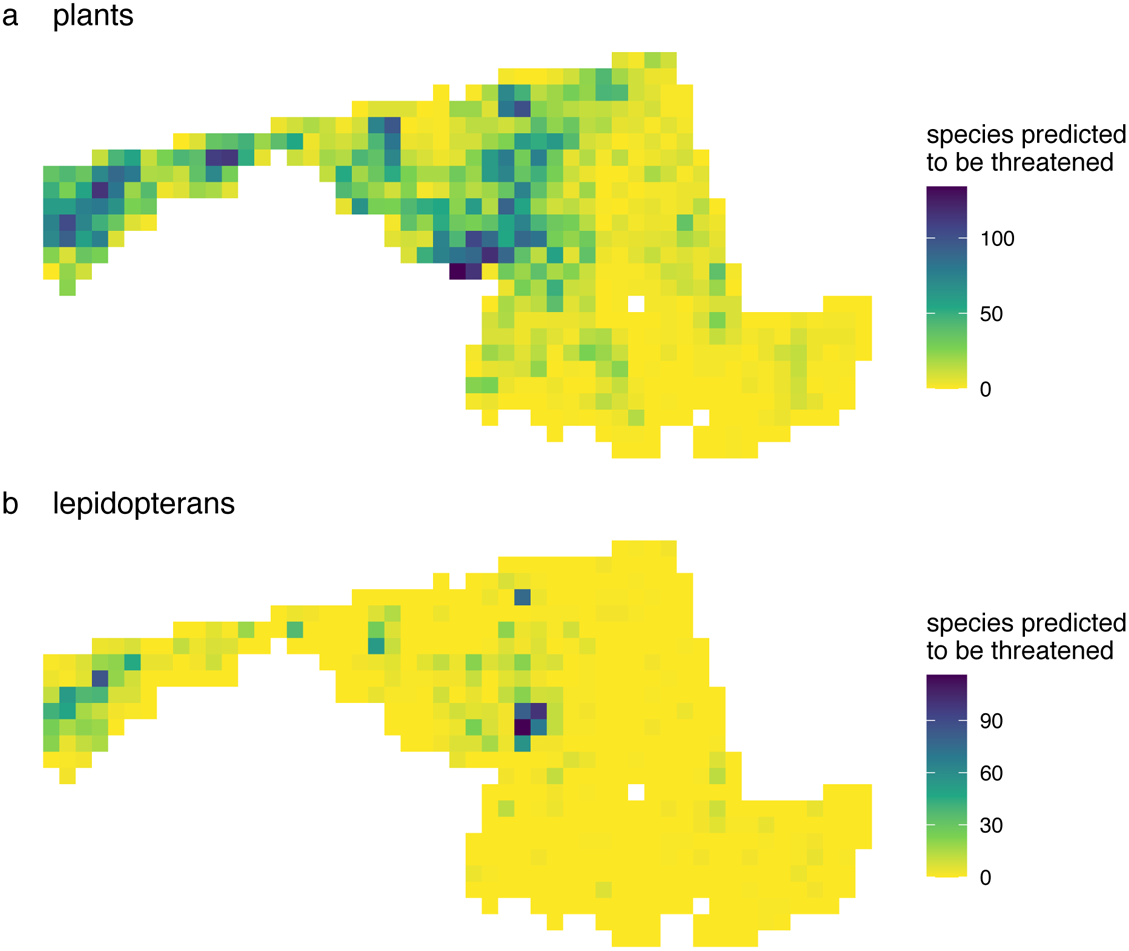
Number of species classified as threatened by the final models using the taxon group-specific probability thresholds (0.66 for plants, 0.36 for lepidoptera), per 8km x 8km grid cell across the state of Maryland. Colors represent number of species (see figure scales).

When corroborating predictions with external data, we found that species with the highest “threatened” probabilities had been assigned a status we considered “threatened” in at least one jurisdiction in eastern North America (20/30); species with low threatened probabilities were rarely unassessed in eastern North America (3/93), and were assigned “threatened” statuses at lower rates (33/93) Recent occurrence patterns for many of the species strongly predicted as “threatened” suggest they have a range primarily north of Maryland, often with southern range extensions along the Appalachian Mountains (Table S1). Lepidopterans receiving the highest “secure” probabilities were less frequently assigned statuses we considered “threatened” in nearby jurisdictions (19/78). The case was different for plants, where 14/15 species had been assigned a “threatened” status elsewhere in eastern North America; primarily to the west and north of Maryland. We note that many of these plant species had also been assessed as relatively secure in coastal states from Delaware southwards, where climates tend to be warmer and where their ranges primarily extend. These patterns also support, to some degree, our predictions.

## Discussion

Here, we showed that random forest models can help cover the Rabinowitzian shortfall, using the biogeographic information about species contained in their occurrence records to evaluate conservation status. We showed that this machine learning method, originally developed for global analysis, applies at regional scales (here, the State of Maryland) and to a variety of taxa. We identified unassessed plant and lepidopteran species that are likely to be threatened and thus could be given high conservation priority. Our prediction mapping highlighted the western, montane section of the state as a potential hotspot for species monitoring, assessment, and protection. In the following sections, we discuss the implications of our results, explore the biogeographic trends recognized by our models, and describe how our study supports the use of similar methods to cover Rabinowitzian shortfalls in additional contexts.

### Publicly Available Data and Simple Predictive Models Identify Regional-Scale Conservation Priorities

Several studies have demonstrated proof-of-concept for predictive modelling approaches for monitoring or conservation prioritization for plant and animal groups, but at continental or global scales (Bastin et al., 2019; Parsons, Pelletier, Wieringa, Duckett, & Carstens, 2022; Pelletier et al., 2018; Wieringa, 2022; Zhang, Campomizzi, & Lebrun-Southcott, 2022; Zizka et al., 2022). Because decisions about monitoring, research, and protection are often made at smaller spatial scales, before recommending adoption of these tools for conservation practice, it is crucial to determine if they still provide useful predictions at those scales (McIntosh et al., 2018; Meyer & Pebesma, 2021; Pressey et al., 2013; Wyborn & Evans, 2021), as we have done here.

We anticipated several obstacles to prediction would be larger at the smaller geographical scale considered here, but also recognized mitigating advantages. Formally similar models trained on global vs. regional datasets may reflect fundamentally different biogeographic processes (e.g., continental vs. local), and are built on data of different quality and quantity. Disadvantages at regional scales include the fact that occurrence records are unlikely to cover complete species’ ranges, thereby obscuring their ecological preferences (A. Lee-Yaw, L. McCune, Pironon, & N. Sheth, 2022; Phillips et al., 2009). Another limitation is that the variation of environmental covariates used is likely smaller, theoretically limiting the strength of any ecological signals and increasing the chance that model predictions would be based on noise (Garcia-Rosello, Gonzalez-Dacosta, Guisande, & Lobo, 2023; Hughes et al., 2021). Finally, and most simply, the number of taxa available on which to train models at regional scales is likely to be much smaller than at global scales, further reducing the chance that predictive models reflect generally applicable trends.

Mitigating some of these concerns, at smaller scales, geographic datasets may have higher resolution and higher quality, or be more consistently available across species or space, which should improve the performance of resulting models. In this vein, we used complete, publicly-accessible, high-resolution covariates at the regional scale, such as land use, land cover, and their history, as well as slope – information that may be less readily available at much larger scales (Gogol-Prokurat, 2011). Though subject to many of the environmental and spatial gaps and biases typical among large biodiversity data (Bachman, Nic Lughadha, & Rivers, 2018; Gallagher et al., 2023; Kitzes & Shirley, 2016; Newbold, 2010), occurrence datasets from our study region benefit from high densities of monetary wealth, human population, and roads, as well as proximity to flagship natural history collections and older research universities. These advantages may, in this study, have counterbalanced the limitations of the smaller spatial scale discussed above.

Overall, even as our regional datasets differ from those used in larger scale predictive models, they appear to contain sufficient information to generate credible predictions. First, we found evidence that the models were using stable – and perhaps robust – signals to generate predictions. Across cross-validation runs of models for the same higher taxon, the same top predictors were frequently identified, implying sample sizes and signal strengths were adequate within our regional-scale data. At the same time, we note some signs of model instability: performance differences in the inner and outer cross-validation nesting levels for plants, as well as high variability in hyperparameter selection especially for lepidopterans. Overall, however, the resulting models discriminated well between previously assessed threatened and relatively secure species. We found binary thresholds for each higher taxon at which previously assessed species were classified accurately (albeit at quite different vote counts; 36% for lepidopterans and 66% for plants). Relatively low representation of threatened species among assessed lepidopterans (25%) versus plants (70%) may partly explain differences in the selected thresholds (Peirce, 1884).

Our models showed strong performance on unseen data via cross-validation. Nevertheless, because we do not suspect that the training data (assessed species) are a representative sample of all species, we remain cautious in our claims of success. We know *a priori* that species have historically been prioritized for assessment based on a variety of social and ecological factors that, while potentially shared among assessed species, may not predict threat across other species that have not been historical priorities for monitoring or protection (Bachman et al., 2019; Brummitt et al., 2015; Di Marco et al., 2017; Fraixedas, Roslin, Antão, Poyry, & Laine, 2022; Ricketts, Daily, & Ehrlich, 2002). A strong test of our predictions would require performing formal status assessments across a large, representative subsample of previously unassessed species, which is beyond the scope of the present study. Here, we informally validated a small subset of the predictions using external lines of evidence, which supported prediction credibility, and thereby the use of this predictive approach at such a regional scale.

### Predicted Conservation Needs Reflects Biogeographic Trends

Generally, our predictions received support from the external data sources, with some caveats. Species our models predicted most likely to be threatened were much more likely to have been assigned a “threatened” status elsewhere in eastern North America, and also more likely to have a “threatened” status assigned south of Maryland, despite the fact that fewer taxa had status assessments in the southeastern USA than in the northeast and Canada. Additionally, trends in recent occurrences of species predicted as “threatened” and “secure” with high probabilities provide biogeographic context and support to our predictions.

Our models predicted the likely statuses of species, but their occurrences point to spatial patterns relevant in the context of regional conservation and climate change. First, we found that the species our models predicted as “threatened” were concentrated in the northwestern part of the state, which includes a central portion of the Appalachian Mountains. Second, our models predicted threat in both taxon groups for northern-ranged, likely cold-adapted species, and relative security for southern-ranged, likely warmer-adapted species. Within the State of Maryland, range-edge species with few populations in the State are considered conservation priorities, and our results suggest that for both plants and lepidoptera, cold-adapted, range-limited species may be especially vulnerable. More generally, montane habitats harbor a high proportion of total, rare, and threatened species (Körner, 2004) and the Appalachians are no exception (Jenkins, Van Houtan, Pimm, & Sexton, 2015). The Appalachian Mountains are home to species considered relicts from glacial periods (Pickering, Kays, Meier, Andrew, & Yatskievych, 2003) that face increasing threats from climate warming (La Sorte & Jetz, 2010; Zhu, Papeş, Armsworth, & Giam, 2022). It is thus expected that our predictions would also highlight montane regions of Maryland as potential conservation hotspots. Furthermore, the northwestern half of Maryland contains many habitats recognized as key for wildlife conservation (Maryland Department of Natural Resources, 2015b), as well as priority areas for conservation and monitoring (Maryland Department of Natural Resources, 2015a), constituting a compelling stage for long-term conservation (Lawler et al., 2015). Beyond the Appalachian region, our results recovered a large number (but not proportion) of species predicted as threatened in the Piedmont region of the State, which has seen rapid development over the past 50 years (Maryland Department of Planning, 2010). Sheer densities of roads, institutions, money, and people in the center of the state may explain this result. However, that section of Maryland is experiencing stark habitat loss and fragmentation, and may also be an emerging priority for monitoring, status assessment, and conservation.

Even as they are not designed for inference, our models recovered ecological signals within (and between) the plant and lepidopteran models in measures of variable importance. These signals point towards mechanisms that drive threat for some species and relative security for others. low variation in mean daily temperature range across occurrence records was a strong predictor of threat for both lepidopterans and plants. Otherwise, top predictor variables differed between the taxon groups, and likely reflect the ecology of each group. For plants, additional variables associated with niche breadth, such as the variation in mean temperature during the coldest month and during the growing period were important, along with land use history and growing season length. Niche breadth has long been associated with vulnerability (Slatyer, Hirst, & Sexton, 2013), and plants restricted to microclimates or habitat types that are rare in Maryland are likely under threat from anthropogenic change. For lepidopterans, the top variables reflected spatial limitations (maximum latitude, minimum longitude, and mean Maryland latitude), as well as number of days below 5°C, all of which we interpret to suggest that more cold-associated lepidopterans tend to be more threatened in Maryland. Abundance and range shifts and declines in response to changing climate are well-documented in lepidopterans (Crossley et al., 2022; Hill, Kawahara, Daniels, Bateman, & Scheffers, 2021), and it is likely that some of Maryland’s most threatened insect fauna are retreating to relatively cool and high-elevation refugia.

### Applications

Our methodology is not intended to replace species conservation assessments (Zizka et al., 2022) nor to stand in for highly contextual, place-based ecological knowledge (Wyborn & Evans, 2021), but rather as a tool that can inform prioritization in a resource-limited conservation world (Sinclair et al., 2018). As resources for protection are already biased towards monitoring and research, approaches like ours can use existing data to accelerate movement of conservation resources towards active conservation work (Buxton et al., 2020). Maryland’s 2015-2025 State Wildlife Action Plan calls for greater coherence in monitoring efforts, especially for species that have yet to be recognized as “of greatest conservation need” (Maryland Department of Natural Resources, 2015c). Towards these ends, first, our models provide guidance about assessed and unassessed plant and lepidopteran species (Table S1) in a coherent, quantitative framework that can immediately integrate new data as they become available. Second, beyond individual species predictions, we also produce regional maps (Fig. 3, S1), that highlight priority areas where conservation needs likely intersect; these maps can support proactive, place-based conservation (Cardillo & Meijaard, 2012). Third, we tentatively identify ecological features of threatened and relatively secure plants and lepidopterans, which warrant further investigation and may be applicable in other regions or to other taxa as well. Our results echo calls to protect climate refugia and unique habitat types within Maryland, and perhaps to assist climate-driven migration (Maryland Department of Natural Resources, 2015a). Finally, as the data inputs for the approach we used are publicly available, as are the scripts containing our workflows (https://datadryad.org/stash/share/cRMHicQVP1f7adWywanzoZ-ftypfUGyXPhPqpQwVaNk), any interested party can reproduce our analysis, update our predictions with new data, or extend our method to additional taxa and locations.

Future applications of our approach would ideally involve stakeholders (Knight et al., 2008). In our study, direct decision-maker input during model construction might have altered which data sources were used, how species were grouped into individual models (here, we generated separate models for plants and lepidopterans), and would also have affected how the models were trained on those data. Decision-makers, uniquely positioned to name the problems associated with different types of misclassification errors, could help translate these problems into loss functions and classifier training criteria (Hand, 2009; Walker et al., 2020). These criteria help tune the model to a specific application (e.g., by avoiding predicting that species are threatened when they are not, or the opposite), via hyperparameter values and vote threshold selections. One advantage of our approach is that it accommodates iterative, stakeholder-driven model construction (Pressey et al., 2013): models were fit on a personal laptop; the decision trees at the heart of random forests are concrete, easily apprehended, and thus accessible; the data we used are publicly available and well-documented.

Overall, we find this modelling approach to be well-suited to use by local and regional conservation planners and decision-makers.

## Conclusion

Predictive methods cannot replace localized knowledge systems nor systematic species assessments (Betts et al., 2020). Instead, in a fast-changing world in which most species, including threatened species, remain unassessed and data-deficient, they extend the conservation utility of existing data. Here, we showed that a predictive method for prioritizing unassessed species, originally developed for global applications, can be flexibly applied at smaller, regional scales and to a variety of taxa. We show how using occurrence records as input data enables spatially explicit predictions, which can highlight priority locations, and could facilitate place-based or ecosystem-level conservation. We highlighted Maryland (USA) unassessed plant and lepidopteran species likely to be threatened, and identified the western part of the state and Appalachian Mountains as hotspots for such likely-threatened species. Such emergent results can help resolve a central conservation mystery: which taxa are most vulnerable to which drivers (Isbell et al., 2023; Sánchez-Bayo & Wyckhuys, 2019). The random forests we used “learn” from existing, publicly accessible datasets using simple decision trees, require relatively little computational resources or natural history expertise to implement, and can be used “in house” by researchers, planners, or managers at regional scales. As a result of the immediacy of our method, as well as the encouraging results from the plant and lepidopteran case studies we present here, we advocate for regional applications of similar predictive tools to better identify conservation priorities. Finally, future applications of these methods could more explicitly integrate stakeholders, as well as consider other niche dimensions (e.g., species interactions or co-occurrences) into both threat assessment and conservation prioritization.

## References

A. Lee-Yaw, J., L. McCune, J., Pironon, S., & N. Sheth, S. (2022). Species distribution models rarely predict the biology of real populations. Ecography, 2022(6), 1–16. 10.1111/ecog.05877

Bachman, S. P., Field, R., Reader, T., Raimondo, D., Donaldson, J., Schatz, G. E., & Lughadha, E. N. (2019). Progress, challenges and opportunities for Red Listing. Biological Conservation, 234(November 2018), 45–55. 10.1016/j.biocon.2019.03.002

Bachman, S. P., Nic Lughadha, E. M., & Rivers, M. C. (2018). Quantifying progress toward a conservation assessment for all plants. Conservation Biology, 32(3), 516–524. 10.1111/cobi.13071

Baker, D. J., Garnett, S. T., O’Connor, J., Ehmke, G., Clarke, R. H., Woinarski, J. C. Z., & McGeoch, M. A. (2019). Conserving the abundance of nonthreatened species. Conservation Biology, 33(2), 319–328. 10.1111/cobi.13197

Bastin, J.-F., Finegold, Y., Garcia, C., Mollicone, D., Rezende, M., Routh, D., … Crowther, T. W. (2019). The global tree restoration potential. Science, 365(6448), 76–79. 10.1126/science.aax0848

Bénard, C., Da Veiga, S., & Scornet, E. (2022). Mean decrease accuracy for random forests: inconsistency, and a practical solution via the Sobol-MDA. Biometrika, 109(4), 881–900. 10.1093/biomet/asac017

Betts, J., Young, R. P., Hilton-Taylor, C., Hoffmann, M., Rodríguez, J. P., Stuart, S. N., & Milner-Gulland, E. J. (2020). A framework for evaluating the impact of the IUCN Red List of threatened species. Conservation Biology, 34(3), 632–643. 10.1111/cobi.13454

Breiman, L. (2001). Random Forest. Machine Learning, (45), 5032.

Brummitt, N. A., Bachman, S. P., Griffiths-Lee, J., Lutz, M., Moat, J. F., Farjon, A., … Lughadha, E. M. N. (2015). Green plants in the red: A baseline global assessment for the IUCN Sampled Red List Index for Plants. PLoS ONE, 10(8), 1–22. 10.1371/journal.pone.0135152

Brun, P., Zimmermann, N. E., Hari, C., Pellissier, L., & Karger, D. N. (2022). Global climate-related predictors at kilometre resolution for the past and future Earth System Science Data Discussions. Earth System Science Data, 14(June), 5573–5603. Retrieved from 10.16904/envidat.332,

Buxton, R. T., Avery-Gomm, S., Lin, H. Y., Smith, P. A., Cooke, S. J., & Bennett, J. R. (2020). Half of resources in threatened species conservation plans are allocated to research and monitoring. Nature Communications, 11(1), 1–8. 10.1038/s41467-020-18486-6

Caetano, G. H. O., Chapple, D. G., Grenyer, R., Raz, T., Rosenblatt, J., Tingley, R., … Roll, U. (2022). Automated assessment reveals extinction risk of reptiles is widely underestimated across space and phylogeny. BioRxiv. 10.1101/2022.01.19.477028

Callaghan, C. T., Nakagawa, S., & Cornwell, W. K. (2021). Global abundance estimates for 9,700 bird species. Proceedings of the National Academy of Sciences of the United States of America, 118(21), 1–10. 10.1073/pnas.2023170118

Cardillo, M., & Meijaard, E. (2012). Are comparative studies of extinction risk useful for conservation? Trends in Ecology and Evolution, 27(3), 167–171. 10.1016/j.tree.2011.09.013

Cardoso, P., Erwin, T. L., Borges, P. A. V., & New, T. R. (2011a). The seven impediments in invertebrate conservation and how to overcome them. Biological Conservation, 144(11), 2647–2655. 10.1016/j.biocon.2011.07.024

Cardoso, P., Erwin, T. L., Borges, P. A. V, & New, T. R. (2011b). The seven impediments in invertebrate conservation and how to overcome them. Biological Conservation, 144(11), 2647–2655. 10.1016/j.biocon.2011.07.024

Chamberlain, S. (2020). Package ‘natserv.’

Chamberlain, S., Oldoni, D., Barve, V., Desmet, P., Geffert, L., McGlinn, D., … Waller, J. (2022). Package ‘rgbif.’ Retrieved from https://github.com/ropensci/rgbif

Chamberlain, S., Szoecs, E., Foster, Z., Arendsee, Z., Boettiger, C., Ram, K.;, … Grenié, M. (2022). taxize: Taxonomic Information from Around the Web. Retrieved from https://docs.ropensci.org/taxize/ (website),%0Ahttps://github.com/ropensci/taxize (devel), https://taxize.dev%0A(user manual)

Crisfield, V., Blanchet, F. G., & Gravel, D. (2022). How and why species are rare: A mechanistic reappraisal of the Rabinowitz rarity framework. 10.22541/au.166748076.66307375/v1

Crossley, M. S., Meehan, T. D., Moran, M. D., Glassberg, J., Snyder, W. E., & Davis, A. K. (2022). Opposing global change drivers counterbalance trends in breeding North American monarch butterflies, (May), 1–10. 10.1111/gcb.16282

Department of the Interior, U. S. F. & W. S. (1973). Endangered Species Act of 1973, As Amended through the 108th Congress. Endangered Species Act Of 1973, 1–44. Retrieved from papers://a25d78e7-d6e0-4b2b-9b83-8d9ae6ec0592/Paper/p134

Di Marco, M., Chapman, S., Althor, G., Kearney, S., Bensancon, C., Butt, N., … Watson, J. E. M. (2017). Changing trends and persisting biases in three decades of conservation science. Global Ecology and Conservation, 10, 32–42. 10.1016/j.gecco.2017.01.008

Eastern Shore Regional GIS Cooperative. (2021). Statewide 1m slope map. Retrieved May 22, 2023, from https://lidar.geodata.md.gov/imap/rest/services/Statewide/MD_statewide_slope_m/ImageServer

Etten, J. Van, Sumner, M., Cheng, J., Baston, D., Bevan, A., Bivand, R., … Mosher, S. (2023). Package ‘raster.’ Retrieved from https://rspatial.org/raster BugReports

Fraixedas, S., Roslin, T., Antão, L. H., Poyry, J., & Laine, A. L. (2022). Nationally reported metrics can’t adequately guide transformative change in biodiversity policy. Proceedings of the National Academy of Sciences of the United States of America, 119(9), 1–4. 10.1073/pnas.2117299119

Free, C. M., Jensen, O. P., Anderson, S. C., Gutierrez, N. L., Kleisner, K. M., Longo, C., … Walsh, J. C. (2020). Blood from a stone: Performance of catch-only methods in estimating stock biomass status. Fisheries Research, 223(November 2019), 105452. 10.1016/j.fishres.2019.105452

Gallagher, R. V., Allen, S. P., Govaerts, R., Rivers, M. C., Allen, A. P., Keith, D. A., … Adams, V. M. (2023). Global shortfalls in threat assessments for endemic flora by country. Plants People Planet, (February), 1–14. 10.1002/ppp3.10369

Garcia-Rosello, E., Gonzalez-Dacosta, J., Guisande, C., & Lobo, J. M. (2023). GBIF falls short of providing a representative picture of the global distribution of insects. Systematic Entomology, (May 2022), 1–9. 10.1111/syen.12589

GBIF.org. (2024). Occurrence Download. 10.15468/dl.9jrwwd

Gogol-Prokurat, M. (2011). Predicting habitat suitability for rare plants at local spatial scales using a species distribution model. Ecological Applications, 21(1), 33–47. 10.1890/09-1190.1

Hand, D. J. (2009). Measuring classifier performance: A coherent alternative to the area under the ROC curve. Machine Learning, 77(1), 103–123. 10.1007/s10994-009-5119-5

Hastie, T., Tibshirani, R., & Friedman, J. (2017). The elements of statistical learning: data mining, inference, and prediction (2nd ed.). Springer. Retrieved from http://www.springerlink.com/index/D7X7KX6772HQ2135.pdf

Heberling, J. M., Miller, J. T., Noesgaard, D., Weingart, S. B., & Schigel, D. (2021). Data integration enables global biodiversity synthesis. Proceedings of the National Academy of Sciences of the United States of America, 118(6), 1–7. 10.1073/pnas.2018093118

Hill, G. M., Kawahara, A. Y., Daniels, J. C., Bateman, C. C., & Scheffers, B. R. (2021). Climate change effects on animal ecology: butterflies and moths as a case study. Biological Reviews, 96(5), 2113– 2126. 10.1111/brv.12746

Hochkirch, A., Samways, M. J., Gerlach, J., Böhm, M., Williams, P., Cardoso, P., … Dijkstra, K. D. B. (2021). A strategy for the next decade to address data deficiency in neglected biodiversity. Conservation Biology, 35(2), 502–509. 10.1111/cobi.13589

Hughes, A. C., Orr, M. C., Ma, K., Costello, M. J., Waller, J., Provoost, P., … Qiao, H. (2021). Sampling biases shape our view of the natural world. Ecography, 44(9), 1259–1269. 10.1111/ecog.05926

iNaturalist Network. (2022). Observations · iNaturalist. Retrieved May 17, 2023, from https://www.inaturalist.org/observations

Isbell, F., Balvanera, P., Mori, A. S., He, J. S., Bullock, J. M., Regmi, G. R., … Palmer, M. S. (2023). Expert perspectives on global biodiversity loss and its drivers and impacts on people. Frontiers in Ecology and the Environment, 21(2), 94–103. 10.1002/fee.2536

Ishwaran, H. (2007). Variable importance in binary regression trees and forests. Electronic Journal of Statistics, 1, 519–537. 10.1214/07-EJS039

Jenkins, C. N., Van Houtan, K. S., Pimm, S. L., & Sexton, J. O. (2015). US protected lands mismatch biodiversity priorities. Proceedings of the National Academy of Sciences of the United States of America, 112(16), 5081–5086. 10.1073/pnas.1418034112

Kitzes, J., & Shirley, R. (2016). Estimating biodiversity impacts without field surveys: A case study in northern Borneo. Ambio, 45(1), 110–119. 10.1007/s13280-015-0683-3

Knapp, W. M., & Naczi, R. F. C. (2020). Vascular plants of Maryland, USA: A comprehensive account of the state’s botanical diversity. Washington, D. C.: Smithsonian Institution Scholarly Press.

Knight, A. T., Cowling, R. M., Rouget, M., Balmford, A., Lombard, A. T., & Campbell, B. M. (2008). Knowing but not doing: Selecting priority conservation areas and the research-implementation gap. Conservation Biology, 22(3), 610–617. 10.1111/j.1523-1739.2008.00914.x

Körner, C. (2004). Mountain biodiversity, its causes and function. Ambio, 33(SPEC. ISS. 13), 11–17. 10.1007/0044-7447-33.sp13.11

Kuhn, M., Wing, J., Weston, S., Williams, A., Keefer, C., Engelhardt, A., … Hunt, T. (2022). Package ‘caret.’

La Sorte, F. A., & Jetz, W. (2010). Projected range contractions of montane biodiversity under global warming. Proceedings of the Royal Society B: Biological Sciences, 277(1699), 3401–3410. 10.1098/rspb.2010.0612

Lawler, J. J., Ackerly, D. D., Albano, C. M., Anderson, M. G., Dobrowski, S. Z., Gill, J. L., … Weiss, S. B. (2015). The theory behind, and the challenges of, conserving nature’s stage in a time of rapid change. Conservation Biology, 29(3), 618–629. 10.1111/cobi.12505

Liaw, A., Wiener, M., Breiman, L., & Cutler, A. (2022). Package “randomForest.”

Ling, C. X., Huang, J., & Zhang, H. (2003). AUC: A statistically consistent and more discriminating measure than accuracy. IJCAI International Joint Conference on Artificial Intelligence, 519–524.

Maryland Department of Natural Resources. (2015a). Conservation actions. In 2015-2025 State Wildlife Action Plan. Retrieved from https://dnr.maryland.gov/wildlife/Documents/SWAP/SWAP_Chapter7.pdf

Maryland Department of Natural Resources. (2015b). Maryland’s Key Wildlife Habitats. In Maryland State Wildlife Action Plan 2015-2025. Retrieved from https://dnr.maryland.gov/wildlife/Documents/SWAP/SWAP_Chapter4.pdf

Maryland Department of Natural Resources. (2015c). Monitoring and Effectiveness Measures. In Maryland State Wildlife Action Plan 2015-2025.

Maryland Department of Planning. (2010). A summary of land use trends in Maryland. Retrieved from https://planning.maryland.gov/Documents/OurProducts/landuse/MDP2010_LU_Summary.pdf

Maryland Natural Heritage Program. (2021a). List of Rare, Threatened, and Endangered Animals.

Maryland Natural Heritage Program. (2021b). Rare, Threatened, and Endangered Plants of Maryland. Retrieved from http://dnr2.maryland.gov

Massicotte, P., South, A., Hufkens, K., & Philippe, M. (2023). Package ‘rnaturalearth.’ Retrieved from https://docs.ropensci.org/rnaturalearth/

McIntosh, E. J., Chapman, S., Kearney, S. G., Williams, B., Althor, G., Thorn, J. P. R., … Grenyer, R. (2018). Absence of evidence for the conservation outcomes of systematic conservation planning around the globe: A systematic map. Environmental Evidence, 7(1), 1–23. 10.1186/s13750-018-0134-2

Meyer, H., & Pebesma, E. (2021). Predicting into unknown space? Estimating the area of applicability of spatial prediction models. Methods in Ecology and Evolution, 12(9), 1620–1633. 10.1111/2041-210X.13650

Mokany, K., & Ferrier, S. (2011). Predicting impacts of climate change on biodiversity: A role for semi-mechanistic community-level modelling. Diversity and Distributions, 17(2), 374–380. 10.1111/j.1472-4642.2010.00735.x

NatureServe. (2023). NatureServe Explorer. In NatureServe. Retrieved from https://explorer.natureserve.org/

Newbold, T. (2010). Applications and limitations of museum data for conservation and ecology, with particular attention to species distribution models. Progress in Physical Geography, 34(1), 3–22. 10.1177/0309133309355630

Nongame and endangered species conservation act. (2022). Code of Maryland. 10.1101/2023.08.31.555788

Panter, C. T., Clegg, R. L., Moat, J., Bachman, S. P., Klitgård, B. B., & White, R. L. (2020). To clean or not to clean: Cleaning open-source data improves extinction risk assessments for threatened plant species. Conservation Science and Practice, 2(12), 1–14. 10.1111/csp2.311

Parsons, D. J., Pelletier, T. A., Wieringa, J. G., Duckett, D. J., & Carstens, B. C. (2022). Analysis of biodiversity data suggests that mammal species are hidden in predictable places. Proceedings of the National Academy of Sciences of the United States of America, 119(14). 10.1073/pnas.2103400119

Pebesma, E., Bivand, R., Racine, E., Sumner, M., Cook, I., Keitt, T., … Dunnington, D. (2023). Package ‘sf.’ Retrieved from https://r-spatial.github.io/sf/

Peirce, C. S. (1884). The numerical measure of the success of predictions. Science, 4(93), 453–454. 10.1126/science.ns-4.93.453.b

Pelletier, T. A., Carstens, B. C., Tank, D. C., Sullivan, J., & Espíndola, A. (2018). Predicting plant conservation priorities on a global scale. Proceedings of the National Academy of Sciences, 115(51), 13027–13032. 10.1073/pnas.1804098115

Phillips, S. J., Dudík, M., Elith, J., Graham, C. H., Lehmann, A., Leathwick, J., & Ferrier, S. (2009). Sample selection bias and presence-only distribution models: Implications for background and pseudo-absence data. Ecological Applications, 19(1), 181–197. 10.1890/07-2153.1

Pickering, J., Kays, R., Meier, A., Andrew, S., & Yatskievych, K. (2003). The Appalachians. In P. R. Gil, R. A. Mittermeier, C. G. Mittermeier, J. P. Pilgrim, G. Fonseca, W. R. Konstant, & T. Brooks (Eds.), Wilderness Earth’s Last Wild Places (pp. 458–467). Conservation International, Washington, DC. Retrieved from http://www.discoverlife.org/co/

Pressey, R. L., Mills, M., Weeks, R., & Day, J. C. (2013). The plan of the day: Managing the dynamic transition from regional conservation designs to local conservation actions. Biological Conservation, 166, 155–169. 10.1016/j.biocon.2013.06.025

Probst, P., Wright, M. N., & Boulesteix, A. L. (2019). Hyperparameters and tuning strategies for random forest. Wiley Interdisciplinary Reviews: Data Mining and Knowledge Discovery, 9(3), 1–15. 10.1002/widm.1301

Rabinowitz, D. (1981). Seven forms of rarity. The Biological Aspects of Rare Plant Conservation, 205–217. 10.1177/004728702237415

Regan, H. M., Hierl, L. A., Franklin, J., Deutschman, D. H., Schmalbach, H. L., Winchell, C. S., & Johnson, B. S. (2008). Species prioritization for monitoring and management in regional multiple species conservation plans. Diversity and Distributions, 14(3), 462–471. 10.1111/j.1472-4642.2007.00447.x

Ricketts, T. H., Daily, G. C., & Ehrlich, P. R. (2002). Does butterfly diversity predict moth diversity? Testing a popular indicator taxon at local scales. Biological Conservation, 103(3), 361–370. 10.1016/S0006-3207(01)00147-1

Sánchez-Bayo, F., & Wyckhuys, K. A. G. (2019). Worldwide decline of the entomofauna: A review of its drivers. Biological Conservation, 232(January), 8–27. 10.1016/j.biocon.2019.01.020

Sinclair, S. P., Milner-Gulland, E. J., Smith, R. J., McIntosh, E. J., Possingham, H. P., Vercammen, A., & Knight, A. T. (2018). The use, and usefulness, of spatial conservation prioritizations. Conservation Letters, 11(6), 1–7. 10.1111/conl.12459

Slatyer, R. A., Hirst, M., & Sexton, J. P. (2013). Niche breadth predicts geographical range size: A general ecological pattern. Ecology Letters, 16(8), 1104–1114. 10.1111/ele.12140

Tobias, A., Sander, O., & Beerenwinkel, N. (2022). Package ‘ ROCR.’

Wagner, D. L., Fox, R., Salcido, D. M., & Dyer, L. A. (2021). A window to the world of global insect declines: Moth biodiversity trends are complex and heterogeneous. Proceedings of the National Academy of Sciences of the United States of America, 118(2), 1–8. 10.1073/PNAS.2002549117

Walker, B. E., Leão, T. C. C., Bachman, S. P., Bolam, F. C., & Nic Lughadha, E. (2020). Caution Needed When Predicting Species Threat Status for Conservation Prioritization on a Global Scale. Frontiers in Plant Science, 11(April), 1–4. 10.3389/fpls.2020.00520

Wickham, H., Chang, W., Henry, L., Pedersen, T. L., Takahashi, K., Wilke, C., … Dunnington, D. (2022). ggplot2: Elegant Data Visualisations Using the Grammar of Graphics. R package version 3.4.0.

Wickham, J., Stehman, S. V., Sorenson, D. G., Gass, L., & Dewitz, J. A. (2021). Thematic accuracy assessment of the NLCD 2016 land cover for the conterminous United States. Remote Sensing of Environment, 257, 112357. 10.1016/j.rse.2021.112357

Wieringa, J. G. (2022). Comparing predictions of IUCN Red List categories from machine learning and other methods for bats. Journal of Mammalogy, 103(3), 528–539. 10.1093/jmammal/gyac005

Wyborn, C., & Evans, M. C. (2021). Conservation needs to break free from global priority mapping. Nature Ecology and Evolution, 5(10), 1322–1324. 10.1038/s41559-021-01540-x

Zhang, X., Campomizzi, A. J., & Lebrun-Southcott, Z. M. (2022). Predicting population trends of birds worldwide with big data and machine learning. Ibis, 164(3), 750–770. 10.1111/ibi.13045

Zhu, G., Papeş, M., Armsworth, P. R., & Giam, X. (2022). Climate change vulnerability of terrestrial vertebrates in a major refuge and dispersal corridor in North America. Diversity and Distributions, 28(6), 1227–1241. 10.1111/ddi.13528

Zizka, A., Andermann, T., & Silvestro, D. (2022). IUCNN – Deep learning approaches to approximate species’ extinction risk. Diversity and Distributions, 28(2), 227–241. 10.1111/ddi.13450

