## Supplemental table S2 and figure S1 for "Global conservation prioritization approach provides credible results at a regional scale"

**Table S2.** Top and bottom fifteen (or more, in case of ties) species per taxon group, based on the predicted probability of being threatened. Known conservation status from other jurisdictions in eastern North America is shown, if available. Codes indicate state/province conservation status and the state/province in question. Row coloring indicates observed species range type. Purple: iNaturalist occurrence records appeared concentrated north of Maryland, with a southward extension along the Appalachian Mountains; blue: occurrences concentrated north of Maryland, without an Appalachian extension; orange: occurrences appear concentrated in the Appalachian Mountains; green: occurrences neither concentrated to the north of Maryland nor clustered along the Appalachian Mountains; white: unclear.

| taxon group | predicted status | genus | species | vote fraction (threatened) | status elsewhere | notes |
|---|---|---|---|---|---|---|
| lepidopterans | threatened | *Idia* | *diminuendis* | 0.984 | S2 ON; S3 NH; S4 PA | |
| | | *Hyppa* | *xylinoides* | 0.98 | S4 PA, NB, NS; S5 ON | |
| | | *Olethreutes* | *quadrifidum* | 0.98 | S4 ON | |
| | | *Papaipema* | *pterisii* | 0.974 | SX NJ; SH DE; S3 VA, IN; S4 ON | Eats ferns, prefers sandy soils |
| | | *Hypena* | *edictalis* | 0.972 | S3 VA; S4 ON | very distinctive Appalachian extension. Laportea specialist. Note very close spelling with congener |
| | | *Haploa* | *confusa* | 0.968 | S4 ON | Cynoglossum officinale specialist which has a pronounced northern range with appalachian extension |
| | | *Crambus* | *bidens* | 0.966 | S4 ON | |
| | | *Plusia* | *contexta* | 0.966 | S4 ON | |
| | | *Clepsis* | *persicana* | 0.964 | S4 PA, ON | very distinctive Appalachian extension. Host generalist |
| | | *Hedya* | *chionosema* | 0.964 | S4 PA, ON | |

| | | | | | | |
|---|---|---|---|---|---|---|
| | | *Protodeltote* | *albidula* | 0.958 | S2 VA; S4 PA, NB, NS; S5 ON | |
| | | *Schinia* | *florida* | 0.958 | S3 VA; S4 PA | oenothera specialist |
| | | *Sicya* | *macularia* | 0.958 | S4 PA, ON, NB | |
| | | *Nola* | *cilicoides* | 0.956 | S4 ON | |
| | | *Peridea* | *basitriens* | 0.956 | S4 PA, ON, NB | |
| | | *Acronicta* | *impressa* | 0.95 | S4 PA, NB, NS; S5 ON | |
| | | *Acronicta* | *americana* | 0 | S4 NB, NS; S5 PA, ON, PE | |
| | | *Actias* | *luna* | 0 | S4 QE, NB, NS; S5 IN, PA, VT, ON | |
| | | *Agnorisma* | *badinodis* | 0 | SH QE; S4 PA, ON | |
| | | *Agrotis* | *ipsilon* | 0 | S5 PA | |
| | | *Alypia* | *octomaculata* | 0 | S5 PA, ON | |
| | | *Amphipyra* | *pyramidoides* | 0 | S4 ON, NB, NS; S5 PA | |
| | | *Anageshna* | *primordialis* | 0 | S4 PA, ON | |
| | | *Antheraea* | *polyphemus* | 0 | S3 VT; S4 NB, NS; S5 PA, IN, ON | |
| | | *Apantesis* | *phalerata* | 0 | S3 QE; S4 PA, ON | |
| | | *Argyrotaenia* | *velutinana* | 0 | S4 PA; S5 ON | |
| | | *Athetis* | *tarda* | 0 | S3 ON, S4 PA | |
| | | *Atteva* | *punctella* | 0 | S4 ON; S5 PA | |
| | | *Battaristis* | *vittella* | 0 | S4 ON | |
| | **secure** | *Callima* | *argenticinctella* | 0 | S4 ON, PA | |
| | | *Cameraria* | *guttifinitella* | 0 | | |
| | | *Cisseps* | *fulvicollis* | 0 | S4 NB; S5 PA, ON | |
| | | *Cisthene* | *plumbea* | 0 | S3 PA; S4 NJ | |
| | | *Costaconvexa* | *centrostrigaria* | 0 | S5 PA, ON | |
| | | *Crambus* | *agitatellus* | 0 | S4 ON, S5 PA | |
| | | *Crocidophora* | *tuberculalis* | 0 | S4 PA, ON | |
| | | *Cyaniris* | *neglecta* | 0 | S4 ME, QE; S5 NC, TN, VA, KY, DE, WV, IN, PA, CT, VT, ON | Genus seems to be Celastrina |
| | | *Cyclophora* | *packardi* | 0 | S4 ON | |
| | | *Cycnia* | *tenera* | 0 | S4 ON, S5 PA | |
| | | *Digrammia* | *ocellinata* | 0 | S4 PA, ON | |
| | | *Dryocampa* | *rubicunda* | 0 | S4 NB, NS; S5 IN, PA, On | |
| | | *Eacles* | *imperialis* | 0 | SX ME, NH; SH RI, CT; S1 MA, S3 QE; S4 ON; S5 NJ, PA, IN | |

| | | | | | |
|---|---|---|---|---|---|
| | | *Elaphria* | *grata* | 0 | S3 ON, S4 PA |
| | | *Epimecis* | *hortaria* | 0 | S2 ON; S5 PA |
| | | *Epipaschia* | *superatalis* | 0 | S4 PA |
| | | *Euchaetes* | *egle* | 0 | S4 ON; S5 PA |
| | | *Eudonia* | *heterosalis* | 0 | S4 PA, ON |
| | | *Eumorpha* | *pandorus* | 0 | S3 KY; S4 IN, PA; S5 ON |
| | | *Eutrapela* | *clemataria* | 0 | S4 NB; S5 PA, ON |
| | | *Galgula* | *partita* | 0 | S4 PA; S5 ON |
| | | *Haematopis* | *grataria* | 0 | S4 ON; S5 PA |
| | | *Halysidota* | *harrisii* | 0 | S1 ON; S4 PA |
| | | *Halysidota* | *tessellaris* | 0 | S4 NB; S5 PA, ON |
| | | *Haploa* | *clymene* | 0 | SH QE; S3 ON; S5 PA |
| | | *Helicoverpa* | *zea* | 0 | S4 PA, NB; S5 ON |
| | | *Hemaris* | *diffinis* | 0 | S4 ON, S5 PA, KY, IN |
| | | *Hemaris* | *thysbe* | 0 | S4 KY, NB, NS; S5 IN, PA, ON |
| | | *Hyalophora* | *cecropia* | 0 | S3 VT; S4 IN, ON; S5 PA |
| | | *Hypena* | *scabra* | 0 | S4 NB; S5 PA, ON |
| | | *Hypercompe* | *scribonia* | 0 | S3 ON; S5 PA |
| | | *Hyphantria* | *cunea* | 0 | S4 NB; S5 PA, ON |
| | | *Hypsopygia* | *olinalis* | 0 | S4 PA, ON |
| | | *Idia* | *aemula* | 0 | S4 ON, NB; S5 PA |
| | | *Idia* | *americalis* | 0 | S4 NB; S5 PA, ON |
| | | *Isa* | *textula* | 0 | S2 QE; S4 PA, ON |
| | | *Lacinipolia* | *renigera* | 0 | S4 NB, NS; S5 PA, ON |
| | | *Lascoria* | *ambigualis* | 0 | S4 ON; S5 PA |
| | | *Malacosoma* | *americana* | 0 | S4 NB, NS; S5 PA, ON |
| | | *Marimatha* | *nigrofimbria* | 0 | S1 NY; S4 PA |
| | | *Mythimna* | *unipuncta* | 0 | S4 NB, NS; S5 PA, ON |
| | | *Nadata* | *gibbosa* | 0 | S4 NB; S5 PA, ON |
| | | *Nephelodes* | *minians* | 0 | S4 NB, NS; S5 PA, ON |
| | | *Nomophila* | *nearctica* | 0 | S5 PA, ON |
| | | *Orgyia* | *leucostigma* | 0 | S5 PA, ON |
| | | *Orthonama* | *obstipata* | 0 | S4 NB; S5 PA, ON |
| | | *Palthis* | *angulalis* | 0 | S4 NB; S5 PA, ON |

| | | | | | | |
|---|---|---|---|---|---|---|
| | | *Palthis* | *asopialis* | 0 | S3 ON; S4 PA | |
| | | *Panopoda* | *rufimargo* | 0 | S3 ON; S5 PA | |
| | | *Parallelia* | *bistriaris* | 0 | S4 ON, NB; S5 PA | |
| | | *Parectopa* | *robiniella* | 0 | | |
| | | *Phalaenostola* | *larentioides* | 0 | S4 PA, ON | |
| | | *Phyllocnistis* | *liriodendronella* | 0 | | |
| | | *Platynota* | *idaeusalis* | 0 | S4 PA; S5 ON | |
| | | *Pleuroprucha* | *insulsaria* | 0 | S4 PA, ON, NB | |
| | | *Prochoerodes* | *lineola* | 0 | S4 NB; S5 PA, ON | |
| | | *Prolimacodes* | *badia* | 0 | SH QE; S4 ON; S4 PA | |
| | | *Protodeltote* | *muscosula* | 0 | S4 NB, NS; S5 PA, ON | |
| | | *Pseudeustrotia* | *carneola* | 0 | S4 NB; S5 PA, ON | |
| | | *Renia* | *adspergillus* | 0 | S5 ON | |
| | | *Schinia* | *arcigera* | 0 | S2 QE; S4 PA, NY, ON | |
| | | *Scopula* | *limboundata* | 0 | S4 NB; S5 PA, ON | |
| | | *Spilosoma* | *virginica* | 0 | S4 NB; S5 PA, ON | |
| | | *Synchlora* | *aerata* | 0 | S4 PA, NB; S5 ON | |
| | | *Zale* | *lunata* | 0 | S4 ON; S5 PA | |
| **plants** | **threatened** | *Physalis* | *pubescens* | 0.954 | S3 NC; S4 FL, KY, VA, DE | Knapp and Naczi list var. integrifolia as a waif but var. pubescens as a MD native |
| | | *Oenothera* | *perennis* | 0.95 | S1 GA; S2 SC, KY; S3 IN, NJ; S4 VA, WV, PE; S5 PA, NY, VT, ON, QE, NB, NS | |
| | | *Bidens* | *vulgata* | 0.948 | S3 NC, QE, NS; S4 DE, NB; S5 IN, KY, VA, WV, PA, NJ, NY, ON, VT | |
| | | *Sphenopholis* | *nitida* | 0.94 | S1 ON, VT; S2 MA; S4 NC, KY, WV, DE, NJ; S5 VA, PA, NY | |
| | | *Carex* | *kraliana* | 0.932 | S1 WV, S4 MS, GA, SC, IN | |
| | | *Poa* | *autumnalis* | 0.932 | SX NJ; S1 PA; S2 DE; S4 NC; S5 SC, WV, VA | |
| | | *Triadenum* | *fraseri* | 0.926 | S1 TN, NC; DE; S2 VA; S3 NJ; S4 WV, PA; S5 NY, VT, ON, QE, NB, NS, PE | |

| | | | | | |
|---|---|---|---|---|---|
| | | *Dendrolycopodium* | *hickeyi* | 0.922 | S1 SC; S2 NC; S3 VA, WV, NJ, IN, PE; S4 KY; ON, QE; NB; NS; S5 NY, VT |
| | | *Cornus* | *alternifolia* | 0.92 | S2 MS, FL; S3 NJ; S4 GA, NC, DE ON, PE; S5 VA, WV, PA, NY, VT, ON, NB, NS |
| | | *Pyrola* | *americana* | 0.92 | SX KY; S2 TN, NC, IN, DE; S4 NJ, VT, ON, QE, NB; S5 VA, WV, NY, NS |
| | | *Calystegia* | *spithamaea* | 0.916 | SH CT; S1 DE, MA; S2 NC, CT, ME; S4 KY, WV, ON |
| | | *Euphorbia* | *commutata* | 0.916 | S1 NC, MI, ON; S2 FL; S3 GA; S4 KY, WV, VA |
| | | *Polygala* | *verticillata* | 0.916 | S1 RI, NB; S2 MA, VT; S3 NC, ON; S4 VA, DE S5 BA, KY, WV, PA |
| | | *Polypodium* | *appalachianum* | 0.916 | S1 ON, PE; S2 SC; S3 QE, NB, NS; S4 NC, WV, VA, NJ; S5 KY, NY, VT |
| | | *Solidago* | *squarrosa* | 0.916 | SH KY, DE; S1 NC, IN, NJ; S2 OH, VT; S3 VA, QE; S4 WV, ON, NB; S5 NY |
| | secure | *Carex* | *comosa* | 0.148 | SH KY; S1 MS, PE; S2 TN, WV, NB, NS; S3 NC, QE; S4 VT; S5 VA, DE, NJ, PA, NY, ON |
| | | *Carex* | *annectens* | 0.156 | S1 QE, NB, PE; S22 ON; S3 NC; S4 DE; S5 KY, WV, VA, PA, NY, VT |
| | | *Andropogon* | *glomeratus* | 0.172 | S1 OH; S3 NY; S4 KY, WV; S5 GA, SC, VA |
| | | *Carex* | *longii* | 0.174 | SH VT, ON; S1 OH, WV; S2 KY, PA, NS; S3 NY; S4 DE, NJ; S5 SC, VA |
| | | *Polygala* | *mariana* | 0.184 | SX NY; S1 TN; S2 NJ; S4 AL, NC; DE; S5 KY, VA |
| | | *Carex* | *swanii* | 0.186 | SX NB; S2 SC, QE; S3 GA, NC, NS; S4 MS, ON; S5 KY, WV, VA, DE, NJ, PA, NY, VT |
| | | *Cyperus* | *retrorsus* | 0.188 | SH PA; S1 NY, MA; S2 KY; S4 FL, DE; S5 MS, GA, SC, NC, VA |

| | | | | | |
|---|---|---|---|---|---|
| | | *Carex* | *lupulina* | 0.19 | S3 NB, NS; S4 MS, FL, VA, VT, QE; S5 SC, NC, IN, KY, WV, DE, NJ, PA, NY, ON |
| | | *Juncus* | *dichotomus* | 0.192 | SH NH; S1 OH, WV, PA; S2 NY; S4 GA, DE; S5 SC, NC, VA, NJ |
| | | *Lobelia* | *puberula* | 0.192 | S1 PA, S4 KY; S5 SC, NC, VA, WV |
| | | *Hydrocotyle* | *umbellata* | 0.194 | SH PA; S1 OH, CT; S2 NY, NS; S4 NJ; S5 FL, NC, VA, DE |
| | | *Dichanthelium* | *scoparium* | 0.196 | SH WV, S1 IN, OH, PA, NY, MA; S3 FL; S4 NJ; S5 SC, NC, VA, DE |
| | | *Fimbristylis* | *autumnalis* | 0.196 | S1 VT, NS; S2 ME; S4 NC, ON, QE; S5 SC, KY, WV, VA, DE, NJ, NY |
| | | *Phoradendron* | *leucarpum* | 0.196 | SX PA, NY; S3 IN, NJ; S4 OH, WV, DE; S5 FL, NC, VA |
| | | *Elymus* | *virginicus* | 0.198 | S4 WV; S5 NC, KY, IN, NY, ON |

plants

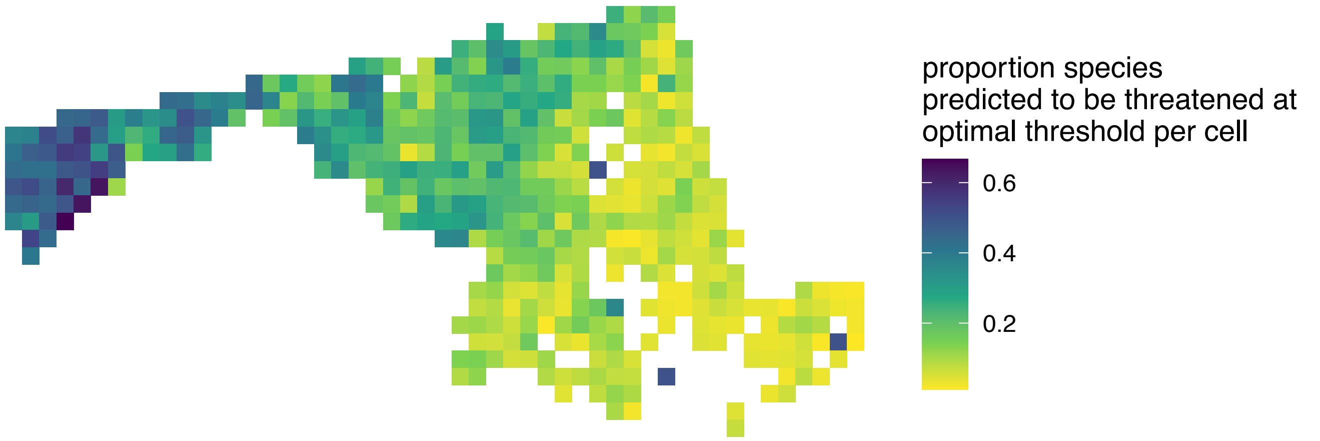

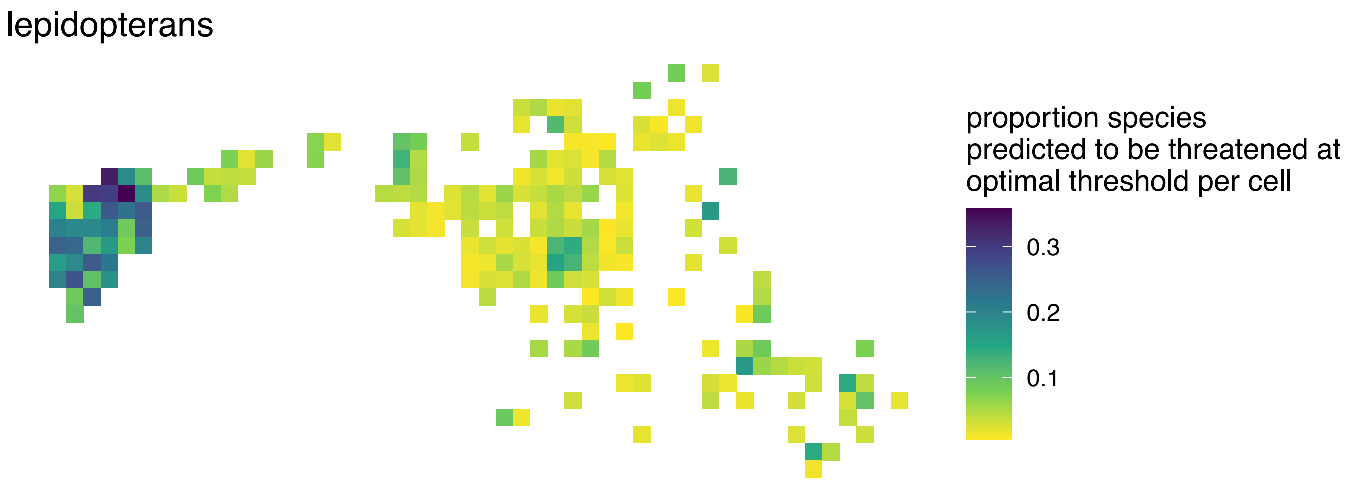

**Figure S1**. The proportion of species predicted as "threatened" in final models using the taxon-group-specific probability thresholds (0.66 for plants, 0.36 for lepidopterans) was highest in western Maryland for both taxa. Proportions computed for each 8km x 8km grid cell across the state of Maryland. Colors represent proportion of species in that category occurring in each cell.
